# *FLOWERING LOCUS T* genes *MtFTb1 and MtFTb2* act redundantly to promote flowering under long days in Medicago truncatula

**DOI:** 10.64898/2025.12.15.694442

**Authors:** Soledad Perez Santangelo, Richard Macknight

## Abstract

Flowering time is a crucial agronomic trait affecting yield and plant performance in different climates and photoperiods. While the molecular pathways regulating photoperiodic flowering are well-characterised in *Arabidopsis* and cereals, they remain poorly understood in many other groups of plants. *FLOWERING LOCUS T (FT)* genes are key components regulating the transition from vegetative to reproductive stages. In the legume *Medicago truncatula*, a forage crop, six FT orthologs are present. To date, only the role of *MtFTa1* has been described, promoting flowering after prolonged exposure to cold (vernalization), while the other five remains unclear. Here we show that *MtFTb1* and *MtFTb2* together are essential for flowering under long-day (LD) photoperiods. Using CRISPR/Cas9, we generated *Mtftb1* single mutants and *Mtftb1/2* double mutants. *MtFTb1* and *MtFTb2* genes act redundantly and are required to up-regulate *MtFTa1* under vernalised LD conditions. While *Mtftb1* mutants flower normally under short-day (SD) and LD conditions, *Mtftb1/2* mutants are delayed specifically under LD but retain a vernalization response. Transcriptomic analysis of wild-type and *Mtftb1/2* mutants shows *MtFTb1/2* are essential for MADS-box gene upregulation and modulate known and candidate flowering regulators under LD. These findings advance our understanding of flowering time control in legumes and may inform strategies to improve forage crop productivity.

**Highlight:** *MtFTb1* and *MtFTb2* genes control flowering under long days *in Medicago truncatula*, revealing clade-specific functions of *MtFT* genes and providing novel insights into the environmental regulation of flowering time in legumes.

## Introduction

Precise coordination of flowering time with environmental cues plays a crucial role in ensuring reproductive success, overall yield, and productivity of crops (Andres and Coupland, 2012). The legume family (Fabaceae) stands out as one of the largest and most economically significant flowering families, ranking second only to cereals (Weller and Ortega, 2015). *Medicago truncatula* is a small annual forage plant used as a model organism in legume research*. M. truncatula* is closely related to several agricultural important temperate legume crops, including *Medicago sativa* (alfalfa), several *Trifolium* species (clovers), *Cicer arietinum* (chickpeas), *Lens culinaris* (lentils), and *Pisum sativum* (peas) (Gepts et al., 2005). In addition to its significance in research, *M. truncatula* is used to feed livestock in Mediterranean countries and Australia due to its high protein content and palatability (Young and Udvardi, 2009). To ensure successful growth and improve crop yields, it is crucial to have a deeper understanding of how flowering time is regulated in *M. truncatula* and other temperate legume crops.

Our current detailed understanding of the genetic mechanisms controlling flowering time comes from *Arabidopsis thaliana* (*A. thaliana*), the small annual brassica model plant species (Andres and Coupland, 2012). At least five distinct pathways play a role in transducing external and internal signals to regulate the timing of flowering. These include the autonomous, ambient temperature, vernalisation (long exposure to cold), hormone, and photoperiod pathways. The signals from these different pathways are integrated by floral integrator genes, where *FLOWERING LOCUS T* (*FT*) plays a central role as a major integrator (Pin and Nilsson, 2012). FT belongs to a protein family characterised by the presence of a phosphatidylethanolamine-binding protein (PEBP) domain. In *A. thaliana*, *AtFT* is expressed in leaves, and the AtFT protein subsequently translocates through the phloem to the shoot apex. There, it interacts with the bZIP transcription factor FLOWERING LOCUS D (FD) and the scaffold protein 14-3-3, forming a complex known as the florigen activation complex (FAC). This complex activates MADS-box genes such as *SUPPRESSOR OF OVEREXPRESSION OF CONSTANS 1 (SOC1)* and the floral meristem identity genes *APETALA1 (AP1)* and *FRUITFULL (FUL)* (Blümel et al., 2015; Kinoshita et al., 2021), playing a crucial role in the transition from the vegetative to the reproductive stage.

*AtFT* transcription is tightly controlled by both positive and negative regulators. In response to inductive long days (LD), *AtFT* transcription is activated by the B-box-containing transcription factor *CONSTANS*, which occurs when the day length exceeds a specific threshold, typically during the longer days of spring (Yanovsky and Kay, 2002). In winter annual *A. thaliana*, *FLOWERING LOCUS C (FLC)*, a key floral repressor, prevents flowering by repressing *AtFT* expression in leaves. After vernalization occurs, *AtFLC* is downregulated, releasing the repression of *AtFT* expression and allowing the initiation of flowering in spring (Michaels and Amasino, 1999; Sheldon et al., 1999). Similar to the winter annual *A. thaliana*, *M. truncatula*, is an LD annual plant, where the photoperiod pathway promotes flowering in spring after vernalisation (Putterill et al., 2013). However, *M. truncatula* lacks the repressor gene *FLC*, and although many *CO-like* orthologs have been identified, their role as activators of flowering time is not conserved (Wong et al., 2014).

In a range of crop species, the number of *FT-like* genes is found to be three to four times higher than in *A. thaliana*, with diverse expression patterns and with roles not only in the control of flowering time, showing greater complexity of their function (Cai et al., 2020; Lee et al., 2013; Putterill and Varkonyi-Gasic, 2016; Zhang et al., 2021).

In legumes*, FT-like* genes are divided into three distinct clades: *FTa*, *FTb*, and *FTc* (Weller and Ortega, 2015). In *M. truncatula*, these clades are represented by six genes*: MtFTa1, MtFTa2, MtFTa3, MtFTb1, MtFTb2,* and *MtFT*c. Interestingly, the late-flowering phenotype of *A. thaliana ft-1* can be rescued by the overexpression of *MtFTa1* and partially by *MtFTb1* and *MtFTc* (Laurie et al., 2011). Both *MtFTa1* and *MtFTa2* are strongly induced by vernalisation followed by warm LD, with no detectable expression in non-vernalised LD conditions. *MtFTa2* expression is particularly responsive to SD conditions, with moderate expression in LD conditions. In contrast, *MtFTa1* shows higher expression levels under LD compared to SD conditions (Laurie et al., 2011). The overexpression of the *MtFTa1* gene in *M. truncatula* results in an early-flowering phenotype. In contrast, the *Mtfta1* mutant flowers later than the wild type (WT) in both SD and LD photoperiods (Laurie et al., 2011). Nevertheless, it flowers earlier in LD compared to SD, retaining its response to photoperiod (Jaudal et al., 2019; Laurie et al., 2011). This suggests that additional factors may contribute to flowering under LD photoperiods.

In the last decade, significant progress has been made in understanding the key players involved in both the negative and positive regulation of flowering time in *M. truncatula*. Repressive mechanisms include a *VERNALISATION2-LIKE VEFS-box* gene (*MtVRN2*). In *A. thaliana AtVRN2* is part of the Polycomb complex involved in the repression of *AtFLC*. In *M. truncatula, Mtvrn2* mutants show premature *MtFTa1* expression, leading to early flowering under vernalised long-day (VLD) conditions (Jaudal et al., 2016). On the other hand, positive regulators include MADS-box floral integrators, such as *MtSOC1* and *MtFUL* genes. It was observed that *MtFULb* accelerates flowering when overexpressed in *A. thaliana* WT plants (Jaudal et al., 2015) and that *MtFUL* expression is altered in *Mtfta1* mutants and *MtFTa1_ox*. *Mtsoc1a* mutants exhibit delayed flowering and short stems, whereas *Mtsoc1a/b/c* triple mutants fail to flower entirely (Fudge et al., 2018; Jaudal et al., 2018; Poulet et al., 2024). Lastly, *M. truncatula MtFDa* and *MtFTa1* were found to work synergistically in the shoot apex to promote flowering, were *MtfdaMtfta1* double mutant completely blocks floral transition (Cheng et al., 2021).

While it is well-established that *M. truncatula MtFTa1* plays a crucial role in the floral transition, particularly under VLD conditions, other genes involved in integrating environmental signals, such as photoperiod information, remain unknown.

In *M. truncatula*, the closely related genes *MtFTb1* and *MtFTb2* are located in tandem on chromosome 7 and share 94% protein sequence identity. These genes are primarily expressed in leaves and show minimal expression in other tissues. Photoperiod strongly influences the expression of *MtFTb1* and *MtFTb2*, suggesting their potential contribution to the promotion of flowering under LD conditions (Laurie et al., 2011; Thomson et al., 2021). *MtFTb1* displays a bimodal expression pattern with peaks at 4 hours after dawn (ZT4) and at dusk (ZT16) under LD conditions, whereas *MtFTb2* only exhibits a peak at ZT4 and lacks a peak at dusk (Perez-Santangelo et al., 2022). Considering the expression pattern and responses to day-length conditions and the similarity between *MtFTb1* and *MtFTb2*, we hypothesise that they both function in the regulation of flowering time under LD in *Medicago truncatula*.

In this study, we used CRISPR/Cas9 to generate *MtFTb1* single and *MtFTb1/2* double mutants to investigate their roles in photoperiodic flowering. The *Mtftb1* single mutants were phenotypically similar to wild type under both LD and SD conditions. In contrast, the *Mtftb1/2* double mutants displayed a delayed flowering phenotype under LD but not under SD conditions, while retaining a normal response to vernalisation. Molecular characterization revealed that *MtFTb1* and *MtFTb2* act redundantly to upregulate *MtFTa1* under vernalised LD (VLD) conditions. Transcriptomic profiling of *Mtftb1/2* mutants further revealed their role in regulating both known and candidate flowering time regulators in response to photoperiod.

## MATERIALS AND METHODS

### Vector Construction

The construct for CRISPR-Cas9-mediated mutagenesis was done following the protocol by (Curtin et al., 2017) with the *Glycine max* ubiquitin (*Gmubi*) promoter-driven CRISPR/Cas9 cassette and *Arabidopsis thaliana* AtU6 driven gRNA. Suitable targets within the *MtFTb1* or both the *MtFTb1* and *MtFTb2* genes were identified using the online tool CRISPR2 (http://crispr.hzau.edu.cn/CRISPR2/). To generate the single gene-edited *Mtftb1* lines, one target (sgRNA1) binding only to the *MtFTb1* gene at the end of exon 1 was used. To generate the double gene-edited *Mtftb1/2* mutants, we designed sgRNA1.1, which targets both *MtFTb1* and *MtFTb2* at the end of exon 1. The sgRNA guide sequences can be found in Table S4.

### Plant Transformation

*Agrobacterium tumefaciens*-mediated transformation of *Medicago truncatula var.* R108 was performed using a slightly modified version of the established protocol (Cosson et al., 2006). In brief, leaf tissue explants were incubated with *the A. tumefaciens strain EAH105,* which was transformed with the CRISPR-Cas9 construct. After co-cultivating the leaf tissue with *A. tumefaciens* for 3 days, explants were transferred to media for callus induction with the selecting agent, BASTA and incubated in the dark, with tri-weekly transfers. After approximately 6 weeks, callus tissue was transferred to shoot induction media under a 16-hour light/8-hour dark photoperiod at 24°C. Developing shoots were transferred to rooting media to encourage root development before being planted in soil. T0 lines were selfed to obtain T1 lines, which were genotyped using target-specific PCR and Sanger sequencing. The binary vector pSC218GG contains 35S::BAR cassette for phosphinothricin (BASTA) herbicide selection of transgenic plants.

### Growth conditions

Scarified seeds (seed surface scratched using fine sandpaper) were water imbibed and germinated at room temperature in a Petri dish in the dark. When radicles extended to a sufficient length (∼10 mm), the seedlings were transferred into growth media (3 parts soil (Yates Professional Potting Mix, Auckland, New Zealand) and 1 part vermiculite) in controlled environment rooms. Plants were grown in a growth cabinet supplied with white light (TL5 28W low-pressure mercury discharge lamp, Philips, Auckland, New Zealand) ∼90-110 µmol/m−2s−1) in 65% relative humidity at 25 °C or in a growth room, supplied with white light (∼80-100 µmol/m−2s−1) in ∼60% relative humidity at 18-23 °C. For vernalised conditioning, Medicago seeds were scarified and water imbibed overnight in the dark at room temperature before being placed at 4 °C in the dark for two weeks in a Petri dish. The seedlings were then planted into soil and put into the growth cabinet under the specified light conditions.

### Genomic DNA Extraction and Genotyping

Genomic DNA was extracted from young leaf tissue using the CTAB protocol. PCR amplification was performed using EmeraldAmp® GT PCR Master Mix with target-specific primers. The PCR conditions were as follows: initial denaturation at 98 °C for 5 min, 35 cycles of denaturation at 98 °C for 15 s, annealing and extension at 60 °C for 15 s, and final extension at 72 °C for 20s/kb. Amplification products were analysed by agarose gel electrophoresis, purified using the DNA Clean & Concentrate Kit (Zymo), and sent for sequence analysis (Macrogen, www.macrogen.com). Data were analysed using Geneious®. Primer sequences are listed in Table S4.

### Flowering Time Analysis

For the analysis of flowering time, plants were grown under two different photoperiods: long days (16 hours of light and 8 hours of darkness) and short days (8 hours of light and 16 hours of darkness), that have been previously vernalised or non-vernalised for two weeks, all at a constant temperature of 22 °C. Flowering time was determined by counting the number of nodes produced to the appearance of the first flower in the primary axis.

### cDNA Synthesis and Quantitative RT-PCR

Total RNA was extracted using the Spectrum Plant Total RNA kit (Sigma-Aldrich®). DNAse I (Thermo Scientific®) was used following the manufacturer’s instructions, to remove any contaminating gDNA. Genomic DNA-free RNA was then used to synthesize first-strand cDNA with oligodT using SuperScript™ III Reverse Transcriptase (Thermo Scientific®). Synthesized cDNA (diluted 1 in 10) was used to analyse the expression of genes using real-time quantitative RT-PCR (qPCR). The Medicago SERINE/THREONINE-PROTEIN PHOSPHATASE 2A (PP2A; also known as PROTODERMAL FACTOR 2; *MtPDF2*: Medtr6g084690) genes was used as the internal control (Kakar et al., 2008). cDNA was amplified with qPCRBIO SyGreen Mix (PCR Biosystems) using the LightCycler 480 (Roche). All reactions were carried out following the manufacturer’s protocol, and negative controls for each primer pair were included by replacing the template with nuclease-free ddH2O. The cycling conditions were as follows: initial denaturation at 95℃ for five minutes, followed by 50 cycles of denaturation at 95 °C for 10 s, primer annealing at 58 ℃ for five seconds, and extension at 72℃ for eight seconds. Relative gene expression levels were calculated using the 2(−delta delta C(T)) method (Bookout and Mangelsdorf, 2003). Primers used for qRT-PCR experiments can be found in Table S4.

### RNA-seq Analysis

Total RNA was extracted using the Spectrum Plant Total RNA kit (Sigma-Aldrich, St. Louis, MO) from fully expanded leaf tissue of gene-edited *Mtftb1/2* and control plants grown under LD (16h light/8h dark) for 14 days after 2 weeks of vernalization (Vernalised conditions) or plants grown for 60 days under LD (16h light/8h dark) without previous vernalization (non-vernalised conditions). Three biological replicates were used. Library construction and sequencing were performed at Otago Genomics Facility (OGF) (https://www.otago.ac.nz/genomics) (Illumina NextSeq 2000, 100 bp paired-end reads, providing about 18M unique reads/sample or 36M PE reads/sample).

Sequence reads were mapped to the *Medicago truncatula* reference genome *(Medtr 4.0v2)* (Tang et al., 2014) using HISAT2 (Kim et al., 2019). Before differential expression analysis, genes with fewer than 10 reads on average per condition were discarded (minRDs >= 0.05). Differential gene expression was estimated using the ASpli package version 2.0.0 (Mancini et al., 2021), which leverages the statistical framework implemented in the edgeR R-package (Robinson et al., 2010) to assess statistically significant changes in gene expression. Resulting p-values are adjusted using a false discovery rate (FDR) criterion (Benjamini and Hochberg, 1995). Genes were considered differentially expressed genes (DEGs) if they had an FDR value lower than 0.05 and a log2 fold change (FC) equal or greater than 0.58 (FC >=1.5), or equal or lower than −0.58 (FC <= −1.5). The list of DEGs of the *Mtftb1/2* gene-edited mutant compared to control plants for both growing conditions can be found in Supplementary Data Sets S1 and S2. Hierarchical clustering heatmaps were performed in iDEP: integrated Differential Expression & Pathway analysis online portal (http://bioinformatics.sdstate.edu/idep/). Gene ontology was performed using g:Profiler (https://biit.cs.ut.ee/gprofiler/) (Peterson et al., 2019). The list of potential flowering genes, used for comparison with the DEGs of the *Mtftb1/2* gene-edited mutant, was obtained from Thomson et al. (2019) and expanded by identifying *M. truncatula* genes with Arabidopsis homologs annotated as flowering-related genes.

### Statistical analysis

Data are presented as mean ± standard error of the mean (SEM) or median and interquartile range (IQR). Pairwise comparisons between genotypes WT(R108) vs either *Mtftb1/2* or *Mtftb1* within each condition (NVLD, VLD, NVSD, VSD) were done using Wilcoxon test. Statistical significance was determined at α = 0.05, with significance levels indicated as follows: *p < 0.05, **p < 0.01, ***p < 0.001, ****p<0.0001. All statistical analyses were performed using R software (https://www.r-project.org/).

## RESULTS

### Loss of function mutation in *MtFtb1* does not alter flowering time

We set out to examine the role Medicago *MtFTb* genes play in regulating flowering. Previous studies have shown that the knockout of *MtFTb2* (due to a *Tnt1* insertion) did not significantly affect flowering time (Thomson et al., 2021). We therefore investigated the involvement of the related gene, *MtFTb1,* in the regulation of photoperiodic control of flowering. Unfortunately, no Tnt1 insertion has been identified within the early coding region of the *MtFTb1* gene. Therefore, we used CRISPR-Cas9 gene editing to generate mutations in the *MtFTb* genes.

*MtFTb1* and *MtFTb2* share 95% nucleotide identity over their coding regions (Laurie et al., 2006; Figure S1). To generate mutations in the *MtFTb1* gene but not in *MtFTb2*, we specifically designed a guide RNA (sgRNA1) targeting a region of *MtFTb1* exon 1 with nucleotide mismatches between the two *FTb* genes (Figure 1A, Figure S1). Following a self-cross, we performed target-specific PCR and Sanger sequencing to confirm the desired edits. Mutations in the *MtFTb1* gene were identified in a single line. The *Mtftb1-1* mutant line had a homozygous insertion of a single A base within exon 1 (Figure 1A; Table S1). The insertion caused a frameshift, resulting in a truncated protein with the first 61 of 173 amino acids, followed by seven new amino acids before terminating with a premature stop codon (Figure S2A).

**Figure 1.**
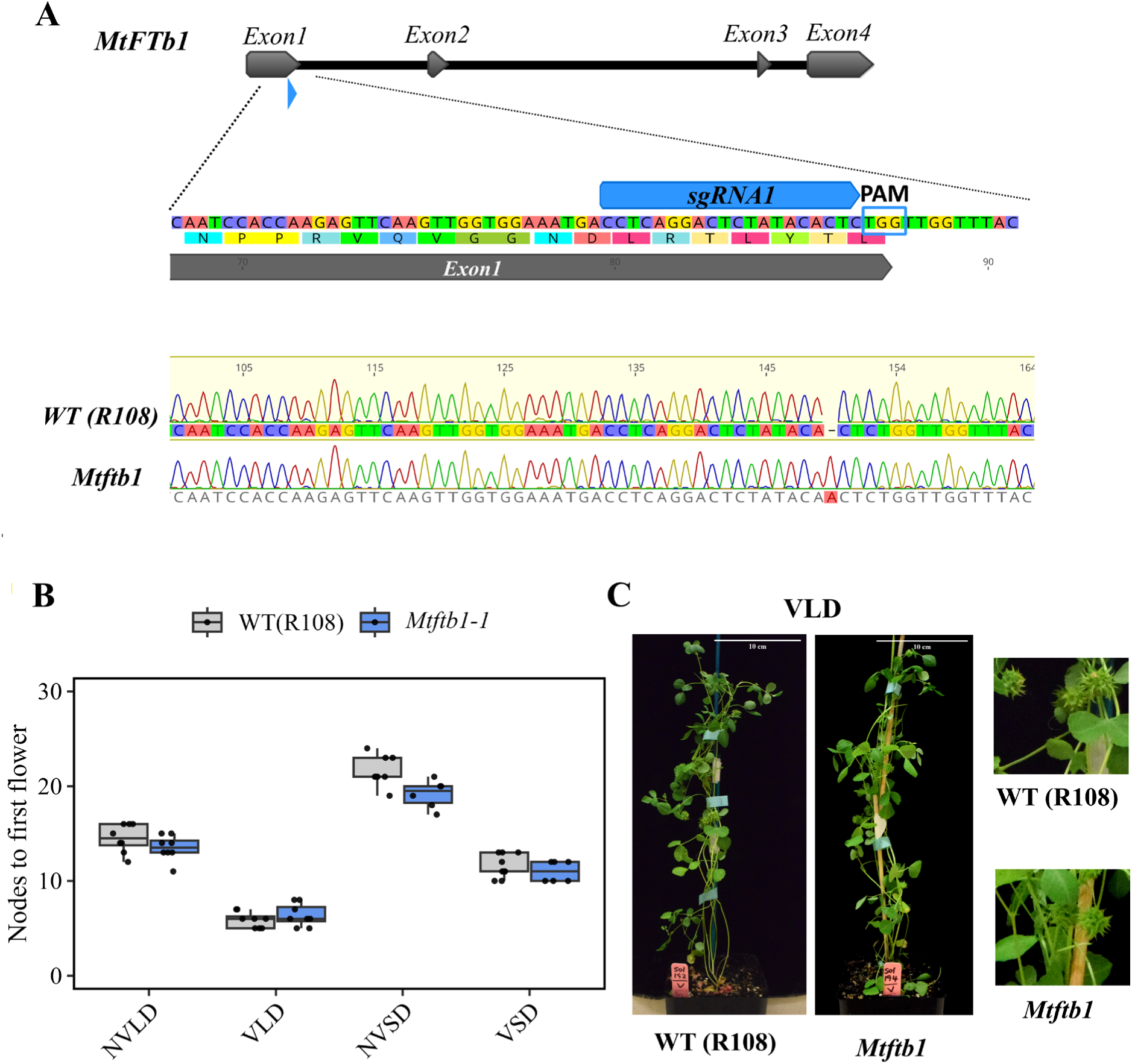
*Mtftb1* gene-edited mutant does not affect flowering time. **A**) Gene schematics indicating the sgRNA1 binding site in *MtFTb1* gene, the PAM sequence and Sanger sequencing for both wild-type (WT) (R108) and *Mtftb1-1* mutant line. **B**) Flowering time represented as nodes to the first flower under four photoperiods: NVLD (Non-vernalised Long Day), VLD (Vernalised Long Day), NVSD (Non-Vernalised Short Day), VSD (Vernalised Short Day). **C)** Representative image of 45-day-old WT (R108) and *Mtftb1-1* gene-edited mutant under VLD.

To investigate the contribution of the *MtFTb1* gene to the regulation of flowering time, the homozygous *Mtftb1-1* mutant plants were grown under four different photoperiod conditions: non-vernalised long days (NVLD), vernalised long days (VLD), non-vernalised short days (NVSD) and vernalised short days (VSD). The *Mtftb*1 mutant plants developed normally and exhibited no changes in flowering time compared to the wild-type WT (R108) plants in any of the four growing conditions (Figure 1B-C). This result suggests that *MtFTb1* alone does not significantly impact the regulation of flowering time.

### *Mtftb1/2* gene-edited mutant lines flower later under a long-day photoperiod

Given that both *MtFTb1* and *MtFTb2* are upregulated under long-day photoperiods and encode similar proteins, we next explored the idea that they might function redundantly to control flowering in response to day length. For this, we generated an *Mtftb1/2* double mutant using gene editing, designing a guide, sgRNA1.1, that targets exon 1 of both *MtFTb1* and *MtFTb2* (Figure 2A; Figure S1). As done with the *Mtftb1* mutant, following a self-cross we performed target-specific PCR and Sanger sequencing. We obtained three independent lines with edits in both the *MtFTb1* and *MtFTb2* genes, which we named *Mtftb1/2-1*, *Mtftb1/2-2*, and *Mtftb1/2-3.* Mutations led to single nucleotide deletions or insertions (Figure 2A; Figure S4A; Table S1) that resulted in a frameshift and premature stop codon, creating truncated *MtFTb1* and *MtFTb2* proteins (Figure S2B). We selected the line *Mtftb1/2-1* to perform the detailed physiological and molecular characterisations.

**Figure 2.**
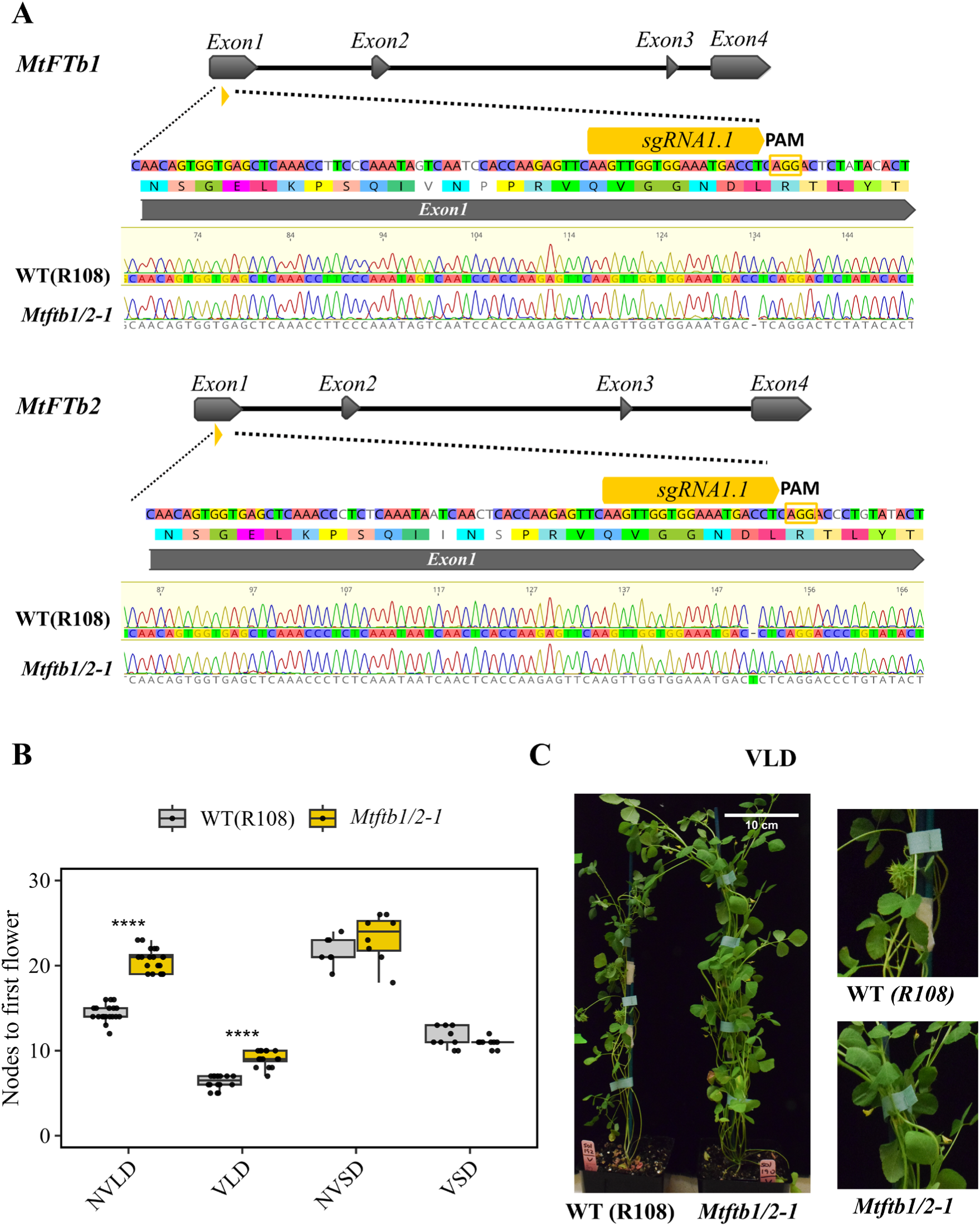
*Mtftb1/2-1* gene-edited mutants have delayed flowering only under inductive long-day photoperiods. **A**) Gene schematics indicating the sgRNA1.1 binding site in both *MtFTb1* and *MtFTb2* genes, the PAM sequence and Sanger sequencing for both wild-type (WT) (R108) and *Mtftb1/2-1* mutant. **B**) Flowering time represented as nodes to the first flower under four photoperiods: NVLD (Non-vernalised Long Day), VLD (Vernalised Long Day), NVSD (Non-vernalised Short Day), VSD (Vernalised Short Day) (Wilcoxon test, ****p<0.0001). **C)** Representative images of 45-day-old plants under VLD conditions.

To determine if the *Mtftb1/2* gene-edited double mutants affected flowering, we analysed the *Mtftb1/2-1* under the same four photoperiodic conditions used to analyse the *Mtftb1* single mutant. When grown under short day (SD) conditions, with or without vernalization, the *Mtftb1/2-1* double mutant flowered like WT plants, with no significant difference (Figure 2B). In contrast, *Mtftb1/2-1* mutant plants exhibited a significant delay in flowering time when grown under LD conditions, irrespective of whether the plants had been vernalised or not. Under LD photoperiod (without vernalisation), the *Mtftb1/2-1* double mutant flowered with 20.6 ± 0.3 nodes compared to 14.4 ± 0.2 for the WT, and under VLD photoperiod (with vernalization) the double mutant flowered with 9.0 ± 0.29 nodes compared to 6.3 ± 0.2 nodes for the WT plants (Figure 2B-C; Figure S3). This was also observed for the *Mtftb1/2-2* and *Mtftb1/2-3* under VLD and LD conditions where both lines flowered later than the WT plants (Figure S4B). This result indicates that both *MtFTb1* and *MtFTb2* are necessary for the induction of flowering under inductive long-day (LD) photoperiods.

### *MtFtb1* and *MtFtb2* are required for the induction of *FTa1* in the LD photoperiod

We previously showed that *MtFTa1* is required for *M. truncatula* to flower in response to vernalization with the *Mtfta1* mutant being ‘blind’ to vernalizing treatment, flowering at the same time with or without vernalization (Jaudal et al., 2019; Laurie et al., 2011). In addition to being upregulated by vernalization, the *MtFTa1* gene is upregulated by LD. Interestingly, *MtFTa1* and *MtFTb1/MtFTb2* genes have different expression patterns with *MtFTb1* and *MtFTb2* having a strong diurnal expression whereas *MtFTa1* is expressed at more similar levels throughout the day (Laurie et al., 2011; Perez-Santangelo et al., 2022).

Interestingly, it has been observed that *MtFTb1* and *MtFTb2* gene expression is downregulated in plants with disrupted photoperiodic regulation, such as *PHYA* photoreceptor mutants (Jaudal et al., 2020) and plants overexpressing the transcriptional repressor *MtCDF1_1* (Zhang et al., 2019), with both lines resulting in late flowering phenotype. Nonetheless, *MtFTb1* and *MtFTb2* expression remains unchanged in *Mtfta1* mutants. This suggests that *MtFTa1* and *MtFTb1*/*MtFTb2* respond differently to long-day photoperiods and raises the possibility that *MtFTa1* is regulated by *MtFTb1* and *MtFTb2*.

To evaluate the potential influence of *MtFTb1* and *MtFTb2* on the expression of *MtFTa1*, we analysed the expression of *MtFTa1* in the gene-edited single *Mtftb1* mutant and *Mtftb1/2* double mutants grown under vernalised long-day (VLD) conditions for two weeks and harvested at ZT4, the time point where the peak expression of both *MtFTb1* and *MtFTb2* occurs. First, for the gene-edited single *Mtftb1* mutant, we observed no significant difference in the expression of *MtFTa1* compared to the wild type (Figure 3A). This aligns with the absence of a flowering time phenotype in the *Mtftb1* mutant. In contrast, we observed a strong downregulation of *MtFTa1* expression in the three *Mtftb1/2* double mutant lines (Figure 3A; Figure S5).

**Figure 3.**
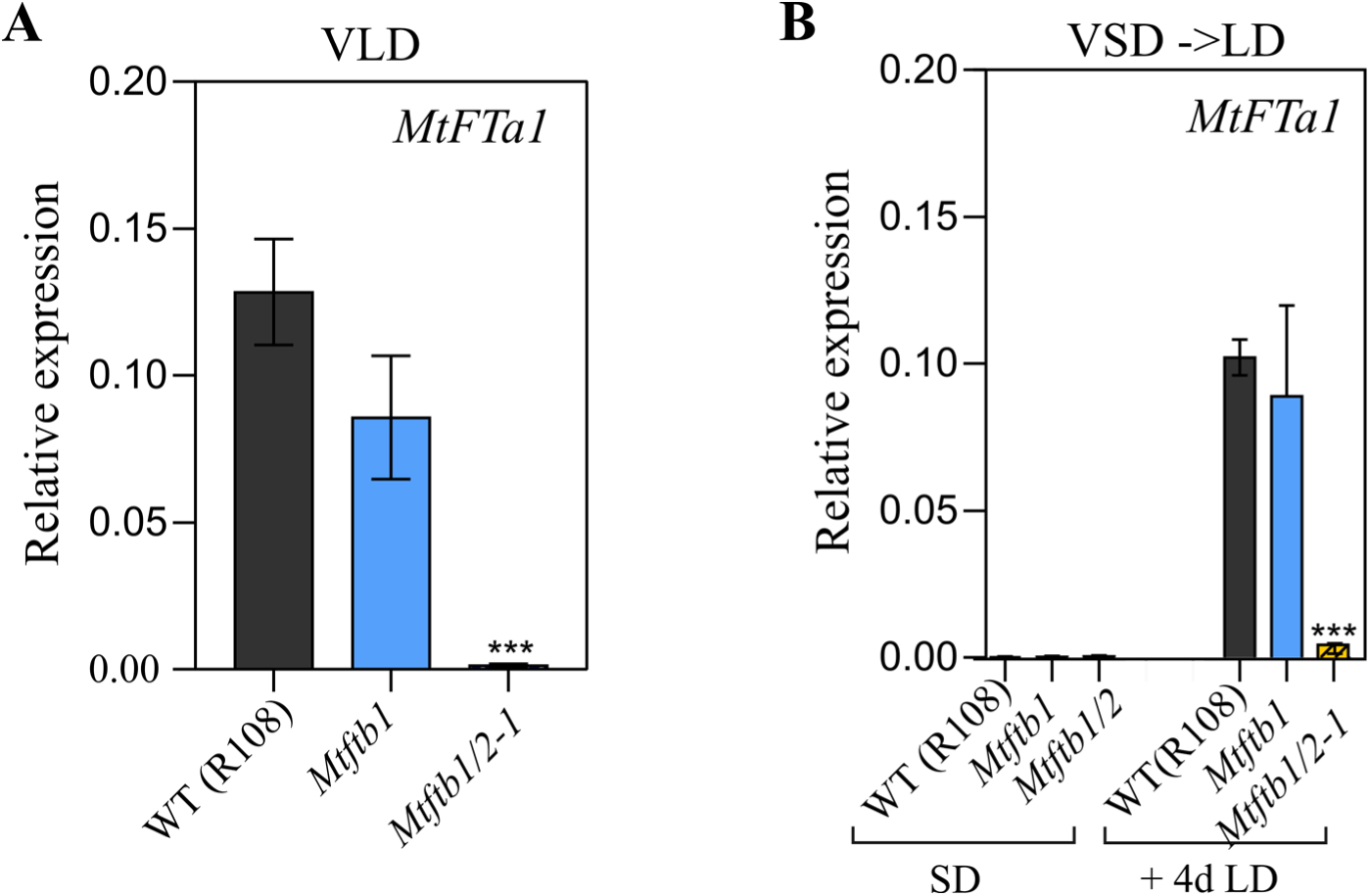
*MtFTa1* is downregulated in the *Mtftb1/2* gene-edited mutant. **A**) Transcript profiles of *MtFTa1* under vernalised long days (VLD). Plants were vernalised for 14 days before being grown for two weeks under long days. The fully expanded youngest trifoliate leaf was harvested at ZT4. **B**) Transcript profiles of *MtFTa1* during the shift from short days (SD) to long days (LDs). Plants were vernalised for 14 days before being grown under SD for 6 days, a group of SD plants was harvested, and the remaining plants were shifted to LD and harvested after 4 days. The fully expanded youngest trifoliate leaf was harvested at ZT4 in both photoperiods. The data represent the average of three biological replicates; each composed of two plants. Relative expression normalized to *MtPDF2* (Medtr6g084690) (Wilcoxon test, ***p<0.001).

Both *MtFTb1* and *MtFTb2*, alongside *MtFTa1*, exhibit a strong response when transitioning from a non-inductive (SD) to an inductive (LD) photoperiod (Laurie et al., 2011). To gain a deeper understanding of the influence of *MtFTb1* and *MtFTb2* on the activation of *MtFTa1*, plants were vernalised for 14 days and grown under non-inductive SD conditions for six days, before transitioning to LD conditions. We collected samples during both SD conditions and four days after the shift to LD conditions at ZT4. Under SD conditions, no expression of *MtFTa1* was detected in both the single *Mtftb1* and double *Mtftb1/2-1* mutant plants, as well as in the WT plants, as expected (Figure 3B). Following the transition from SD to LD conditions, a significant upregulation of *MtFTa1* expression was observed in both the WT and the single *Mtftb1* mutant lines. However, in the double *Mtftb1/2-1* mutants, this upregulation did not occur (Figure 3B).

These findings collectively show that *MtFTa1* expression is induced by *MtFTb1* and *MtFTb2* under long-day (LD) conditions following vernalisation. Linking these two flowering induction mechanisms, photoperiod and vernalisation, ensures the initiation of flowering in *M. truncatula* in spring after the winter cold has passed.

### Identification of differentially expressed genes in *Mtftb1/2* mutants

We found that *MtFTb* genes play a role in the response to LD conditions, both under vernalised and non-vernalised conditions. To explore the regulatory gene network downstream of *MtFTb* genes on a global scale in leaves, we performed RNA sequencing of the gene-edited *Mtftb1/2* vs WT plants. For this, we grew WT and gene-edited *Mtftb1/2* plants under two different conditions: VLD, where the plants were vernalised for 14 days and then grown for two weeks under LD photoperiod, to capture changes in expression profiles in the period in which the plant becomes physiologically committed to flower after the “winter” cold has passed; the second condition was NVLD, where non-vernalised plants were grown for 60 days, a crucial time point when wild-type plants typically undergo the transition to flowering. This approach allowed us to investigate gene expression changes at the flowering time transition without the effects of vernalisation, at a later stage in development. In each condition, we collected wild-type and *Mtftb1/2* leaf samples from three biological replicates at ZT4, capturing the time and tissue where both *MtFTb1* and *MtFTb2* are highly expressed (Laurie et al., 2011; Perez-Santangelo et al., 2022).

A differential gene expression analysis comparing WT vs *Mtftb1/2* VLD samples identified 1164 differentially expressed genes (DEGs), with 650 genes upregulated (FC>=1.5) and 514 genes downregulated (FC<=1.5) in *Mtftb1/2* (SupplementaryDataSet1, Figure 4A). On the other hand, the differential gene expression analysis comparing WT vs *Mtftb1/2* NVLD samples revealed just 246 DEGs, with 160 genes upregulated (FC>=1.5) and 86 genes downregulated (FC<=1.5) in the *Mtftb1/2* mutant (SupplementaryDataSet2, Figure 4B). Only 57 DEGs were shared between the VLD and LD samples, showing that each sampled condition captured distinct gene expression profiles, with *Mtftb1/2* mutation impacting gene expression differently under different developmental stages and environmental conditions (Figure 4C; Supplementary Data Set 3).

**Figure 4.**
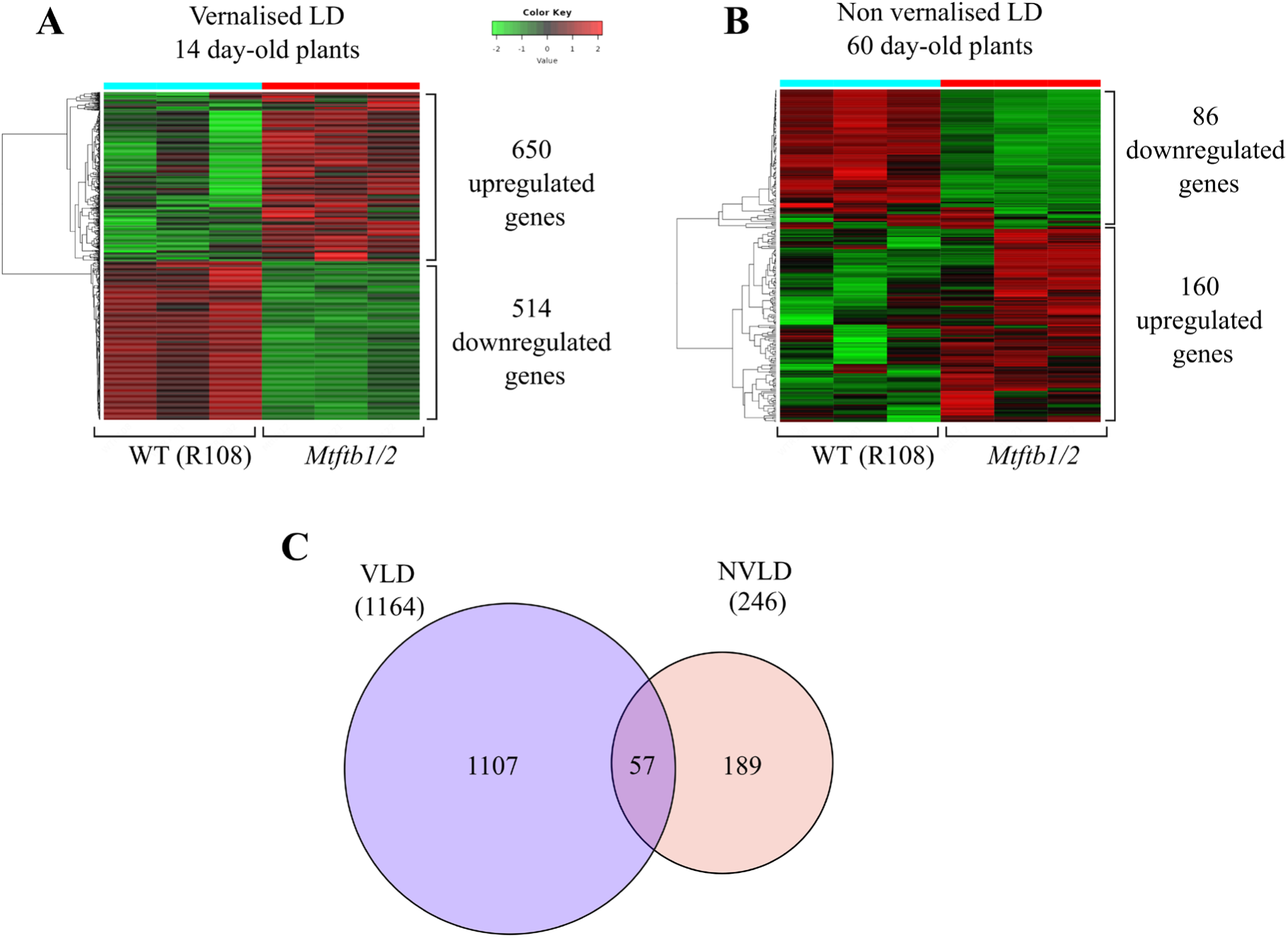
Global analysis of differentially expressed genes in the *Mtftb1/2* gene-edited mutant. **A**) Hierarchical clustering heatmap of up- and down-regulated genes in wild-type (WT) R108 vs. *Mtftb1/2-1* gene-edited mutant for 14-day vernalised plants grown for two weeks under long days (VLD). **B**) Hierarchical clustering heatmap of up- and down-regulated genes in WT vs. *Mtftb1/2-1* gene-edited mutant for 60-day-old plants grown under long days (NVLD). **C**) Venn diagram of shared differentially expressed genes (DEGs) between VLD and NVLD datasets.

### Identification of potential flowering time genes regulated by *MtFTb1/b2*

Interestingly, evidence suggests that *FT* genes in the leaves also play a role in regulating the transcriptional reprogramming of vegetative development, further supporting the transition to flowering (Teper-bamnolker and Samach, 2005). We first used our VLD dataset, conditions where *MtFTa1* is present, to identify potential flowering time genes that might be directly or indirectly regulated by *MtFTb1/MtFTb2* and modulate the expression of *MtFTa1*. Amongst the list, we found *MtE1L* and *MtSPA1L*, to be expressed at lower levels in the *Mtftb1/2* mutant (Table S2). *MtE1L* (Medtr2g058520) is a homologue of an important legume-specific regulator involved in photoperiodic flowering in soybean (Xia et al., 2012). *MtSPA1L* (Medtr8g027985) is a homologue of the *SUPPRESSOR OF PHYA-105* (*SPA*) gene family, which is known to play a role in regulating the key B-box-containing transcription factor, *CONSTANS* (*CO*), and is also involved in light signalling and photomorphogenesis pathways in *A. thaliana* (Laubinger et al., 2006; Sheerin et al., 2015). We also found putative orthologs of *A. thaliana NUCLEAR FACTOR Y (NF-Y) B2* and *B3* (*Medtr4g133938*). *AtNF-YB2/B3* acts as a positive regulator in the photoperiod-mediated flowering time pathways, driving the expression of *FLOWERING LOCUS T* (*FT*) (Takagi et al., 2023). Also, a member of the *SQUAMOSA PROMOTER-BINDING-LIKE* (*SPL*) protein family (*Medtr8g463140*), with homology to miR156 regulated-*AtSPL4*, a key component of the ageing pathway (Table S2). Last, we found a member of the PEBP family, with homology to *BFT (BROTHER OF FT AND TFL1*), which was strongly downregulated (Medtr0020s0120). In *A. thaliana*, *BFT* is known to act as a repressor of flowering time. Given that it was downregulated in the *Mtftb1/2* double mutant, in *M. truncatula,* may play a distinct role as a positive regulator of floral transition, downstream of *MtFTb1/b2* (Table S2).

On the other hand, among the genes expressed at higher levels in the *Mtftb1/2* mutant, we identified two basic helix loop helix (bHLH) transcription factors (Medtr8g062240 & Medtr8g099880), known as *MtCBL10* and *MtCBL9* respectively and Medtr4g070080, a homolog to *AtCCR2* a circadian-regulated gene, also known as *AtGRP7* (Streitner et al., 2008). Last, among the upregulated genes in the *Mtftb1/2* mutant we found a CCT B-box zinc finger protein domain gene (Medtr1g109350) with homology to *AtBBX27*, which does not have a known role in *A. thaliana* (Table S2). Considering the significance CCT B-box zinc finger family of genes in regulating various developmental processes, including the control of flowering time, *MtBBX27-like* (Medtr1g109350) emerges as a potential negative regulator of the photoperiodic flowering pathway.

### *MtFTb1* and *MtFTb2* are required for the upregulation of MADS-box genes in the leaves

When exploring the top downregulated genes in the shared DEG lists for VLD and NVLD (Supplementary Dataset 3; Table S3), three MADS-box transcription factors were among the most strongly downregulated: *MtFULa*(Medtr2g461760), *MtFULb* (Medtr4g109830), and an AtFUL/AtAGL8 ortholog, which we named *MtFUL-like* (Medtr2g461710). GenBank annotation indicates that *MtFUL*-*like* encodes a 62-amino-acid protein that shares almost complete homology with the N-terminal regions of MtFULb and AtFUL proteins. *MtFUL-like* and *MtFULa* are located in close proximity on chromosome 2; *MtFULa* is annotated as a 173-amino-acid protein, whereas *MtFULb*, located on chromosome 4, is annotated as a 232-amino-acid protein. Based on these observations, we believe that *MtFUL-like* (Medtr2g461710) likely is the N-terminal region of *MtFULa* (Medtr2g461760).

It has been shown that *MtFTa1* modulates the expression of *MtFULa* and *MtFULb*, with *MtFULs* transcription levels increasing in plants overexpressing *MtFTa1* (Jaudal et al., 2015; Yeoh et al., 2013). Our data show that the *MtFULa* (*MtFUL-like*) and *MtFULb* genes can be regulated by *MtFTb1/MtFTb2* in the absence of *MtFTa1*, as they are downregulated in the *Mtftb1/2* mutant under NVLD conditions, where *MtFTa1* is not normally expressed, at least at the developmental stage measured (Table S3). Moreover, Medicago *MtFDa* (an interactor of *FT*) is essential for *MtFTa1* activation of the *MtFUL* genes in the apex (Cheng et al., 2021). Our data showed that *MtFDa* was not expressed in the leaves, suggesting that the activation of *MtFUL* genes in the leaves occurs through complex formation with different transcription factors.

When we focused on MADS-box transcription factors downregulated in each condition, four were identified exclusively in NVLD (Table S3). These include, Medtr2g017865 and Medtr8g087860, which are homologous to *AGAMOUS-LIKE 1 (AtAGL1),* known as *MtAGa* and *MtAGb*, respectively; both involved in floral-organ development (Zhu et al., 2018). Medtr4g109810, homologous to *SEPALLATA* (*AtSEP*), known as *MtSEP4,* crucial for floral organ development (Zhu et al., 2018). Last, the flowering integrator, *MtSOC1b (*Medtr8g033250), was also downregulated (Table S3).

On the other hand, amongst the downregulated group of genes solely in VLD conditions, we found Medtr6g464720, a MADS-box transcription factor with similarity to *AtAGAMOUS 4* (*AtAGL4/AtSEP2*), known as *MtAGL6D* (Dreni and Zhang, 2016), and the flowering integrator *MtSOC1a (*Medtr7g075870) (Table S3).

These findings highlight the role of *MtFTb* genes in upregulating MADS-box transcription factors and their importance in regulating flowering time under both NVLD and VLD conditions. However, their potential role in leaves on other developmental processes in leaves cannot be ruled out.

### *MtFTb1/MtFTb2* mediated transcriptional reprogramming of leaves

To gain a more comprehensive understanding of the transcriptional changes resulting from the knockout of the *MtFTb1* and *MtFTb2* genes in leaves, a GO term analysis was done on the DEGs for both conditions (NVLD and VLD). We found a significant overrepresentation of transcripts associated with translation and ribosome biogenesis among the downregulated genes in *Mtftb1/2* under both VLD and NVLD conditions (Supplementary Data Set 4). The activation of *MtFTb1* and *MtFTb2* is likely needed to enhance the translation of transcripts required for reprogramming leaves during the transition from vegetative to reproductive development. We observed terms related to flower development and inflorescence meristem identity in both conditions (NVLD and VLD respectively). Also, downregulated genes only in *Mtftb1/2* VLD plants were overrepresented for terms related to iron import into the cell and oligopeptide transport. These genes are likely involved in the increased transportation needed to support reproductive development and potentially FT transport to the apex (Supplementary Data Set 4).

These findings collectively show the significant and unique role of *MtFTb1* and *MtFTb2* genes in promoting flowering induction under long-day conditions, regardless of whether vernalisation has occurred. The activation of these genes in *M. truncatula* during long-day conditions is mediated by an unknown mechanism, which triggers a cascade of gene expression changes in the leaves. Subsequently, these changes propagate to the apex, initiating the transition from vegetative growth to flowering. Interestingly, transcripts associated with translational activity are downregulated in the *Mtftb1/2* mutant during this transition

## DISCUSSION

The control of flowering, which greatly impacts the reproductive success and yield of many crops, including *M. truncatula*, converges on a key florigen known as FT. In *M. truncatula*, six FT-like genes have been identified, belonging to three distinct clades (a, b, and c) (Laurie et al., 2011). However, the functional relevance of five of the six *MtFT-like* genes remains elusive. In this study, we provide the first evidence that *MtFTb1* and *MtFTb2* play crucial roles in photoperiod-specific responses in *M. truncatula*

### *MtFTb1* and *MtFTb2* function redundantly to control flowering under long-day photoperiods

In our previous work, we showed that *MtFTa1*, one of the six *M. truncatula FT-like* genes has a prominent role in flowering induction in response to vernalisation. The *Mtfta1* mutants show delayed flowering and have a strongly reduced response to vernalisation, yet they retain their photoperiod responsiveness. (Jaudal et al., 2019; Laurie et al., 2011). This raised the question if other *MtFT-like* genes could play a role in photoperiod responsiveness. The expression of *MtFT-like* clade *b* genes are predominantly observed in leaf tissue, with *MtFTb1* and *MtFTb2* being induced under LD conditions, non-responsive to SD and not influenced by vernalisation (Laurie et al., 2011). Given their expression response, we set out to investigate their potential role in the integration of photoperiod information by using CRISPR-Cas9 gene editing to generate *Mtftb1* single and *Mtftb1/2* double mutants. (Figure 1A;2A; S4A). The single gene-edited *Mtftb1* mutant did not affect flowering time under any of the photoperiod conditions measured (Figure 1B). On the contrary, the *Mtftb1/2* gene-edited double mutant lines show a delayed flowering time under LD and VLD conditions (Figure 2B;S3;S4B), but had the same flowering phenotype as the wild-type under either VSD or SD.

Interestingly, the SD legume soybean (*Glycine max*), where flowering is induced under SD photoperiods, two out of ten *FT-like GmFT2a* and *GmFT5a* have been demonstrated to play distinct roles in promoting flowering, with *GmFT2a* being more critical under SD conditions and *GmFT5a* playing a more predominant role under LD conditions (Cai et al., 2020). In *M. truncatula MtFTb1* and *MtFTb2* have been found to control flowering time under inductive LD photoperiod, where *MtFTa1* plays a more dominant role in the induction of flowering after vernalisation has occurred.

Nevertheless, *MtFTb1* and *MtFTb2* are required for the characteristic upregulation of *MtFTa1*, both under VLD conditions and once plants are shifted from VSD to LD and commitment to flowering has occurred (Perrella et al., 2020). This was observed as the characteristic upregulation was absent in the double *Mtftb1/2* mutant plants (Figure 3). Interestingly, in the *Mtfta1* mutant, the expression levels of *MtFTb1* and *MtFTb2* remained unchanged (Laurie et al., 2011). This indicates that *MtFTb1* and *MtFTb2* are upstream of *MtFTa1* and are essential for the integration of photoperiod information under both LD and VLD conditions

### Transcriptome-wide analysis of *Mtftb1/2* mutants in VLD conditions reveals candidate photoperiodic flowering time genes

The photoperiodic control of flowering time has been extensively studied in *A. thaliana*, where *CONSTANS* (*CO*) acts as the central regulator, integrating day-light cues to induce the expression of *AtFT*. While the CO-FT module is present in crop species like rice, legumes have diverged from this pathway, as *CO-like* genes do not seem to be involved in flowering time control (Wong et al., 2014). To identify candidate genes associated with the photoperiodic flowering pathway regulated by *MtFTb1* and *MtFTb2*, we performed a transcriptome-wide analysis of *Mtftb1/2* double mutants vs wild type plants (Figure 4) under NVLD and VLD conditions.

Intriguingly, a homolog of the *A. thaliana BROTHER OF FT AND TFL1 (AtBFT)* gene (*Medtr0020s0120*; *MtBFTL*), a floral repressor of the *FT/TFL1* gene family, was strongly downregulated in the *Mtftb1/2* mutant in VLD conditions. *AtBFT* is found in the leaves, shoot apex, and inflorescence meristem, and the overexpression of *AtTFL1* causes late flowering and floral indeterminacy (Chung et al., 2010; Yoo et al., 2010). Since *MtBFTL* was downregulated in the late-flowering *Mtftb1/2* mutant, it may act as a positive regulator of PEBP family genes, regulated by *MtFTb1/MtFTb2* (Table S2). Although AT5G62040 (*AtBFT*) is the top hit, it also shares homology with AT5G03840 (*TFL1*) and AT2G27550 (*TFL1-like*) and could therefore act as a repressor in *M. truncatula*.

Downregulated genes under VLD conditions also included the legume-specific protein *MtE1L* (*Medtr2g058520*). While the role of *MtE1L* in *M. truncatula* is not fully understood, previous studies suggest its potential as a positive regulator of photoperiodic flowering, as *Medicago e1l* mutants exhibit a slight late-flowering phenotype (Zhang et al., 2016), and photoreceptor *MtphyA* mutants show downregulation of *MtE1L* (Jaudal et al., 2020). Furthermore, the expression of *MtE1L* appears to be diurnally regulated, displaying a temporal expression pattern similar to *MtFTb1*, and its expression is upregulated in leaves during the transition from SD to LD photoperiods (Perez-Santangelo et al., 2022). In *Mtftb1/2* double mutant*, MtE1L* was downregulated, supporting its potential role as a positive regulator of photoperiodic flowering time in *M. truncatula*. However, further research is needed to determine whether it plays a central role in the regulation of photoperiodic flowering.

Another interesting flowering time candidate gene was *MtSPA1L* (Medtr8g027985), which we found to have lower expression in *Mtftb1/2* double mutant (Table S2). In *A. thaliana*, *SPA1* is known to form a complex with *COP1* and play a negative regulatory role in flowering by facilitating the degradation of CO. Therefore, the downregulation of *MtSPA1L* in the late flowering *Mftb1/2* mutant, suggests its potential involvement in the degradation of a flowering repressor.

On the other hand, we observed the upregulation of a B-box zinc finger protein (Medtr1g109350) in *Mtftb1/2* double mutant, which shares a moderate protein identity (38.54%) with *A. thaliana AtBBX27,* which we named *MtBBX27L*. As the soybean *GmBBX27*, *MtBBX27L* only contains two B-box domains without a CCT domain (Shan et al., 2022). While the exact role of *BBX27* is unknown for either species, the *BBX* gene family are known to play crucial roles in regulating several developmental processes, including photoperiodic flowering.

Interestingly, these three candidate photoperiodic flowering time genes, *MtE1L*, *MtSPA1L* and *MtBBX27L* change their expression between SD and LD photoperiods (Thomson et al., 2019), making them strong flowering time candidate genes for further study. Moreover, a comparison of 146 candidate flowering genes described in Thomson et al 2019, responsive to the change in photoperiod conditions (LD vs SD) with the 514 downregulated genes in *Mtftb1/2* double mutant under VLD conditions revealed a statistically significant overrepresentation (Representation factor: 3.4; p < 0.002) (Figure S6; Supplemental DataSet S5). This highlights the impact of *MtFTb1/2* on candidate photoperiodic responsive genes.

### *MtFTb1* and *MtFTb2* together modulate the expression of MADS-box transcription factors in leaves

Through our transcriptomic analysis, we identified several MADS-box transcription factors to be downregulated in *Mtftb1*/2 mutants, including the floral integrator *FUL* and *SOC* genes (Fudge et al., 2018; Jaudal et al., 2018, 2015; Poulet et al., 2024), as well as homologs of *AGAMOUS* and *SEPALLATA* floral identity genes (Table S3), including *MtAGa (*Medtr2g017865*), MtAGb (*Medtr8g087860*), MtAGL6D (*Medtr6g464720) and *MtSEP4 (Medtr4g109810)*. *MtAGa* and *MtSEP4* have also been found to be downregulated in the double mutants for *MtFTa1* and *MtFDa* (an interactor of *FT*) specifically in the apex, which further supports their potential importance in regulating floral development (Cheng et al., 2021).

## Conclusion

Our study provides strong evidence that *MtFTb1* and *MtFTb2* function as key floral integrators within the photoperiodic pathway of *M. truncatula*. Unlike *A. thaliana*, which relies on a single *FT* gene for integrating both vernalisation and photoperiodic cues, *M. truncatula* possesses three distinct *FT* clades - *FTa, FTb*, and *FTc* - reflecting functional diversification following gene duplication. While *MtFTa1* has been established as a central regulator of the vernalisation-induced flowering pathway, our findings reveal that *MtFTb1 and MtFTb2* specifically control flowering in response to long-day photoperiods and act upstream of *MtFTa1*. This functional partitioning highlights the evolutionary divergence of *FT-like* genes in legumes, revealing a more complex regulatory architecture than that observed in model annual Brassica species. By defining the distinct roles of *MtFTb1* and *MtFTb2*, this work advances our understanding of photoperiodic flowering control in temperate legumes and offers valuable insights for the genetic improvement of forage crop productivity.

## Supplementary data

Fig. S1. Coding sequence alignment of *MtFTb1* and *MtFTb2* genes showing the binding sites of sgRNA guides.

Fig. S2. MtFTb protein sequences in *Mtftb1-1* and *Mtftb1/2-1* mutants showing frame shifts.

Fig. S3. Representative pictures of *Mtftb1/2-1* at three flowering stages under VLD.

Fig. S4. Gene edits and flowering-time phenotypes of *Mtftb1/2-2 and Mtftb1/2-3* mutants.

Fig. S5. Transcript profiles of *MtFTa1* under vernalised long days (VLD) for *Mtftb1/2-1 Mtftb1/2-2 and Mtftb1/2-3* mutants.

Fig. S6. Overlap between downregulated genes in *Mtftb1/2-1* mutants under VLD conditions and SD-LD-responsive flowering candidates from Thomson et al., 2019.

Table S1. CRISPR -Cas9 gene edits in *MtFTb1* and *MtFTb2* genes.

Table S2. Potential flowering time genes differentially expressed in *Mtftb1/2-1* mutants

Table S3. MADS-box transcription factors downregulated in *Mtftb1/2-1* mutants

Table S4. List of oligonucleotides used in this study.

## Acknowledgements

This work was supported by grants from the Royal Society of New Zealand Marsden Fund to RCM. SPS was supported by a Marsden Fast Start grant from the Royal Society of New Zealand. The authors wish to thank Dr. Shereen Asha Murugayah for her assistance in the generation of the gene-edited plants. We also thank Professors Jo Putterill and Jim Weller for their helpful comments and critical review of the manuscript.

## Author Contributions

Conceptualization, S.P.S and R.C.M., S.P.S; performed the experiments and data analysis, SPS.; writing - original draft preparation, S.P.S and R.C.M writing - review and editing. All authors have read and agreed to the published version of the manuscript.

## Data Availability

The raw sequencing data have been uploaded to the Biostudies-Array Express collection and are available under accession number E-MTAB-13536.

